# From injury to full repair: nerve regeneration and functional recovery in the common octopus, *Octopus vulgaris*

**DOI:** 10.1101/693390

**Authors:** Pamela Imperadore, Dario Parazzoli, Amanda Oldani, Michael Duebbert, Ansgar Büschges, Graziano Fiorito

**Author notes:** **Corresponding author:** Pamela Imperadore, Association for Cephalopod Research - CephRes, 80133, Napoli, Italy.

## Abstract

Spontaneous nerve regeneration in cephalopod molluscs occurs in a relative short time after injury, achieving functional recovery of the lost capacities. In particular, transection of the pallial nerve in the common octopus (*Octopus vulgaris*) determines loss and subsequent restoring of two functions fundamental for survival, i.e. breathing and skin patterning, the latter involved in communication between animals and concealing. The phenomena occurring after lesion have been investigated in a series of previous studies, but a complete analysis of the changes occurring at the level of the axons and the effects on animals appearance during the whole regenerative process is still missing. Our goal is to determine the course of events following injury. Our goal is to determine the course of events following injury, from impairment to full recovery.

We observed nerve regeneration, end-target re-innervation and functional reconnections between central brain and periphery, using the contralateral nerve in the same animal as internal control. The final architecture of the regenerated nervous tissue does not mirror the original structure, however functionality returns to match the phenotype of an intact octopus, and with no visible impact on the behaviour of the animal. This provides exceptional value to these findings for future studies.

**Summary statement:** Here we report events occurring after interruption of the peripheral neural circuitry in *Octopus vulgaris*, from the dramatic loss of normal functioning to full recovery.

## Introduction

More than 160 years of studies support evidence of outstanding regenerative abilities of cephalopod molluscs, showing that cuttlefish, squid and octopus are all capable of recovering the structure and function of a variety of damaged/lost tissues and parts, including nervous system (review in Imperadore and Fiorito, 2018). As for other organisms, the regenerative capabilities of cephalopods appear to rely on the plasticity expressed at the cellular level, and on the involvement of pluripotent elements such as stem cells or dedifferentiated cells, cell proliferation and migration (Imperadore et al., 2017; Imperadore et al., 2018; Zullo et al., 2017), and to other regulatory and/or trophic factors including the expression of specific genes (for a general review see Birnbaum and Alvarado, 2008; see also Imperadore, 2017).

One of the most remarkable cases of regeneration in cephalopods is represented by the pallial nerve, a paired neural structure connecting central nervous system (i.e. the sub-esophageal mass) to periphery via the stellate ganglia (for description see: Fredericq, 1878; Young, 1929; Young, 1971; Young, 1972). These two long nerves, which extend for around 40 mm in length in an adult octopus of about 250 g body weight having a mantle length of 95 mm, innervate mantle muscles and chromatophores in the skin (see Supplementary Material, **Fig. S1**). Axons of neurons travelling in this nerve are involved in the neural control of respiratory muscles and skin patterning (Imperadore et al., 2017; 2018; Sanders and Young, 1974; Sereni, 1929; Sereni and Young, 1932).

Unilateral traumatic injury of this nerve leads to paralysis of mantle muscles and immediate paling of the skin on the ipsilateral side of the mantle (e.g., Imperadore et al., 2017; Sanders and Young, 1974). The ability of the pallial nerve to regenerate in octopus was discovered by Sereni and Young (1932), who described the phenomenon from a structural and morphological point of view. Subsequently, recovery of the chromatic function of the skin was also reported. In the course of the above mentioned observations, it was shown that degeneration of both stumps occurred within the first 10 days after lesion, followed by an intense regeneration originating from the central stump first and later from the peripheral one (Imperadore et al., 2017; Sereni and Young, 1932; Young, 1972). Hemocytes infiltration was reported to occur for several days after nerve transection at the level of the cut stumps (Sereni and Young, 1932), together with active phagocytosis, scar formation and proliferation (Imperadore et al., 2017). Proliferation and, possibly, re-differentiation of connective tissue cells within the injured nerve, have been first observed by Imperadore and coworkers (2017; 2018).

The first signs of functional recovery of body patterning (Borrelli et al., 2006; Messenger, 2001; i.e. chromatic and textural appearance of the skin; for review see: Packard and Hochberg, 1977) have been reported to occur starting from 30 days post-lesion (Sanders and Young, 1974), but recent studies found that this may occur earlier, i.e. seven days after injury (Imperadore et al., 2017).

Here we integrate previous findings on the morphological and physiological events occurring after severing the pallial nerve in *Octopus vulgaris*, by extending our previous observations after the first 14 days after injury (Imperadore et al., 2017) and reporting a new set of data describing the complete recovery of the nervous components and the corresponding phenotypic appearance. Backfilling of the pallial nerve allowed detection of regenerating fibres in the nerve as early as five days to five months post lesion (p.l.). Physiological recovery, assessed through examination of the conductivity of the axons in the pallial nerve at five months p.l., is here provided. Furthermore, we provided a phenomenological analysis of mantle muscles contraction for breathing and chromatic patterning of the skin from loss of function (after lesion) to full recovery (about one month, up to five months p.l.).

## Materials and Methods

### Animals

A total of 14 *Octopus vulgaris* of both sexes (8♀, 6♂, body weight: 200 - 450 g) were utilized in this study. Animals were caught in the Bay of Naples (Italy) and following the practice available in the GF laboratory at the Stazione Zoologica (Amodio et al., 2014; Fiorito et al., 1990), each animal was identified, sexed, weighed, and housed in an experimental tank with running seawater (60 × 100 × 50 cm). Octopuses were selected for being intact (i.e. no sign of lesion or regenerating parts). Details about animals’ care and laboratory settings are provided in Supplementary Material.

Experiments with live octopuses included in this study were carried out before transposition of Directive 2010/63/EU in Italy (March 2014). Although no authorization was required, all procedures were performed in order to minimize the pain and distress of the animals involved (Andrews et al., 2013; Fiorito et al., 2014; Fiorito et al., 2015; Smith et al., 2013).

### Surgery: nerve transection

*O. vulgaris* were anaesthetized by immersion in 3.5% magnesium chloride hexahydrate (MgCl_2_ 6H_2_O) in sea water for 15 minutes (Grimaldi et al., 2007). Surgery was performed as described in Imperadore et al. (2017). Briefly, octopuses were turned on their ventral side and the right pallial nerve exposed using a skin hooklet and completely transected. To standardize the site of the injury along the nerve, the diameter of stellate ganglion was measured for each animal. The same length computed as distance along the nerve to set the site of the lesion (3.5 mm on average). Left-side pallial nerve remained intact, serving as an internal control.

Animal were returned to their tanks soon after surgery and allowed to recover. Experiments were carried out only on animals that succeeded in recovering their predatory behaviour, and that exhibit a stable and prompt attack behaviour to the live prey (Amodio et al., 2014). See Supplementary Material for care of octopuses after surgery and during experimental observations.

### Analysis of behavioural responses after lesion

From the analysis of video recordings we considered *O. vulgaris*’ predatory performance, and in particular measure of the readiness to attack and of possible asymmetries, i.e. position and orientation (left/right) that the animal assumes during locomotory-predatory responses to attack/approach. We also considered differences in octopus’ body patterning comparing the lesioned and the control sides. In order to explore any other effect of the lesion, we also scored any side-bias in the use of the mantle musculature in providing efficient breathing movements (see Supplementary Material for details: Analysis of animal performances).

### Humane-killing of octopuses

At the selected time-points (five, 30, 45 days and 5 months) octopuses were anaesthetized by immersion in a 3.5% solution of MgCl_2_ in seawater. Animal behavioural responses were also observed and video recorded during anaesthesia; skin reactions to pinch on both sides of the mantle were also assessed using grooved forceps (as described in Butler-Struben et al., 2018) in lesioned and control animals. When terminal anaesthesia was achieved (> 30 min, Grimaldi et al., 2007), death was confirmed by transection of dorsal aorta, in compliance with recommendations included in Annex IV of the Directive 2010/63/EU (see: Andrews et al., 2013; Fiorito et al., 2015). At sacrifice, both nerves were harvested and used for backfilling experiments to evaluate nerve regeneration patterns and target re-innervation.

### Electrophysiology

Only nerves from animals kept alive for five months p.l., prior to undergo backfilling, were used for electrophysiology experiments. Electrical stimulation of axons in the nerve was applied before the injury site with a stimulation electrode and recording of the induced activity was measured close to the stellate ganglion with a recording electrode. Analogous areas were selected for the contralateral control nerves.

Pallial nerve and the stellate ganglion of recovered and control sides of experimental animals were harvested and placed in a saline filled dish. Bipolar hook electrodes (e.g., Büschges et al., 1992; Sauer et al., 1995) were placed at both ends of the pallial nerve and the electrodes were isolated from the surrounding saline by silicon paste (Baysilone, Fa. GE Bayer Silicones). For stimulation of axons at one electrode, rectangular current pulses of 0.2ms duration and variable amplitude were used (generated by a Grass S48 Stimulator via an isolation unit Grass SIU5). The resulting compound spike of axons in the pallial nerve was recorded extracellularly with the second electrode. Signals were pre-amplified by an isolated low-noise preamplifier (model MA101, Electronics workshop, Institute for Zoology, University of Cologne). The signal was further amplified and high- and low-pass filtered (high-pass: 300Hz; low-pass: 3kHz) using a 4-channel amplifier/signal conditioner (model MA102, Electronics workshop, Institute for Zoology, University of Cologne). Signals from the stimulation and recording electrodes were digitized and recorded at a sampling rate of 12kHz using a Micro-1401-3 analog-to-digital converter (Cambridge Electronic Design, Ltd. Cambridge, UK) and Spike2 software (Cambridge Electronic Design). The threshold amplitude of the stimulus for eliciting a compound spike in the pallial nerve was set to 1.0T (Büschges et al., 1992; Hooper and Schmidt, 2017). The amplitudes for the stimuli were normalized to this threshold.

For evaluation of amplitudes of the compound action potentials induced by electrical stimulation of the pallial nerve, for each stimulation strength, the mean and standard deviation were calculated (considering five original compound action potentials per stimulus strength). Mean values of compound action potentials (lesioned and contralateral intact nerve) were also plotted versus stimulus strength.

### Tissue sampling and backfilling

Pallial nerves together with stellate ganglia and surrounding tissues were harvested from sacrificed animals for neural tracing experiments. Backfill protocol was performed following Imperadore et al. (Imperadore et al., 2019). In brief, the far end of the pallial nerve was placed in a Vaseline pool filled with tracer solution (Neurobiotin 5% in distilled water, Vector Laboratories Cat #SP-1120, RRID: AB_2313575) while the remaining tissue was immersed in sea water. The tracer was left to diffuse for around 24 hours at 4°C.

After dye diffusion, the tissues were fixed in 4% PFA in sea water for 1h 30 min at 4°C and later washed in 0.1 M PBS (pH 7.4). Dehydration was performed with ascending ethanol series (20 min each), followed by a mix of ethanol 100% and methyl salicylate (1:1) (20 min) and by two changes of methyl salicylate (five min each). Rehydration was then obtained with descending ethanol series (20 min each). The samples were treated with collagenase/dispase and hyaluronidase in 0.1 M PBS for 30 min at 37° C on shaker (final concentration of the enzymes was 1mg/ml).

After washing in 0.1 M PBS + 1% Triton X-100 (PBS-TX), samples were placed for pre-incubation in 10% NGS in PBS-TX overnight at 4° C and the following day incubated with streptavidin Cy3 conjugated (Jackson ImmunoResearch Cat# JI016160084) (1:200) in PBS-TX + 10% NGS overnight at 4°C. Nuclei counterstain was performed with DAPI (14.3 μM) in PBS-TX at room temperature (2 hour).

Samples were washed in 0.1 M PBS and dehydrated with ascending ethanol series (20 min each). For clearing, samples were immersed in 100% ethanol and methyl salicylate (1:1) for 20 min and then in pure methyl salicylate until the preparation was fully transparent and clear (usually 15 min). For visualization at the microscope, the whole nerve and ganglion were embedded in methyl salicylate in a metal slide (2 or 3 mm thick) closed on both sides by cover slips.

### Imaging of backfilled nerves

Samples were imaged on a Leica TCS SP8X laser scanning confocal microscope or Leica DMi8 microscope using a 10 (zoom factor 0.40) or a 20x air objective (zoom factor 0.75). Tile Z-stack experiments of whole mount tissues were performed on backfilled nerve and stellate ganglion, both for lesioned and control samples. The laser was set at 550 nm for neurobiotin detection, while DAPI was excited at 405 nm. Z-step size was set at 5 μm. Pixel format size was set at 1024×1024.

To visualize connective tissues in the samples an upright multiphoton microscope was used (Leica SP8 MP). Laser Chameleon Vision II (Coherent) was tuned to 1050 nm for simultaneous excitation of SHG and neurobiotin; 405 nm for excitation of DAPI detection. Two non descanned HyDs were utilized to detect SHG signal (filter BP 525/50) and red signal (BP 585/40); one internal HyD was utilized for DAPI objective. In this case 10x (zoom factor 0.40) air objective with a numerical aperture of 0.3 was utilized. Z-stacks with a step size of 10 μm.

Images were processed using Leica software LAS X or Fiji software.

### Experimental design and Statistical Analysis

The number of octopuses used in this study was chosen in the attempt of both, obtaining a statistical representative group and implement the 3Rs strategy, reducing it to its minimum. To further reduce the number of animals used, we obtained behavioural data from previous experiments, conducted in the same laboratory (at Stazione Zoologica, Naples, Italy) in comparable conditions (see Supplementary Material: Animals utilized for statistical analysis). Experiments, data and following analyses were performed following recommendations on experimental design and analysis in laboratory animals (Festing and Altman, 2002) and, more specifically, guidelines for the care and welfare of cephalopods in research (Fiorito et al., 2015). Behavioural observations from video recordings and data analysis were performed by blinded investigators. The experimental design has been assessed through the NC3Rs Experimental Design Assistant (https://www.nc3rs.org.uk/experimental-design-assistant-eda) achieving power > 0.90.

Statistical analyses were performed using SPSS software (SPSS Inc. Released 2009. PASW Statistics for Windows, Version 18.0. Chicago) to analyse *i*. lateralization, *ii*. latency of attack, *iii*. recovery of the breathing function. Asymmetries (i.e. lateralization) and latencies of attack for the 68 octopuses taken into account were noted before anaesthesia/surgery and the day following the procedures. Following Zar (1999) and Siegel and Castellan (1988) we applied McNemar’s test to examine differences in the SIDE animals’ utilized during attack/approach (within groups) for the two conditions (injured and control); comparisons between groups were tested using the Chi-Square test. Yates’s correction for continuity was applied when necessary. Within- and between-groups comparisons on latency of attack data were tested through the Wilcoxon test and Mann-Whitney U, respectively.

For the analysis of the recovery of breathing acts function after regeneration, we applied a Friedman ANOVA followed by Bonferroni correction. Wilcoxon matched-pairs signed-ranks tests were used as *post hoc* procedure.

Whenever appropriate Monte Carlo simulations were utilized to calculate exact probability (Monte Carlo exact module available in SPSS) providing asymptotic p, a Monte Carlo probability value (MCP) and a 95% CI (confidence interval) of the MCP.

In the cases box and whisker plots were utilized to plot data, we used the SPSS graphical output standards (i.e. boxes represented the interquartile range, 25th-75th percentiles, bars within boxes the median values, whiskers the 10th and 90th percentiles, and outliers are marked). In all cases, statistical differences were considered significant at p < 0.05.

## Results

### Effect of the lesion on octopus’ behaviour

After severing the right pallial nerve and following recovery after anaesthesia all *O. vulgaris* returned to their dens, behaved normally and did not exhibit any sign of distress. In some instances and immediately after recovery from anaesthesia, octopuses exhibited grooming manoeuvre (see description in Borrelli et al., 2006) close to the injured area and/or inside the mantle cavity (Fig. 1 E) on the side of lesion. In most of the instances, grooming disappeared after few hours.

**Figure 1.**
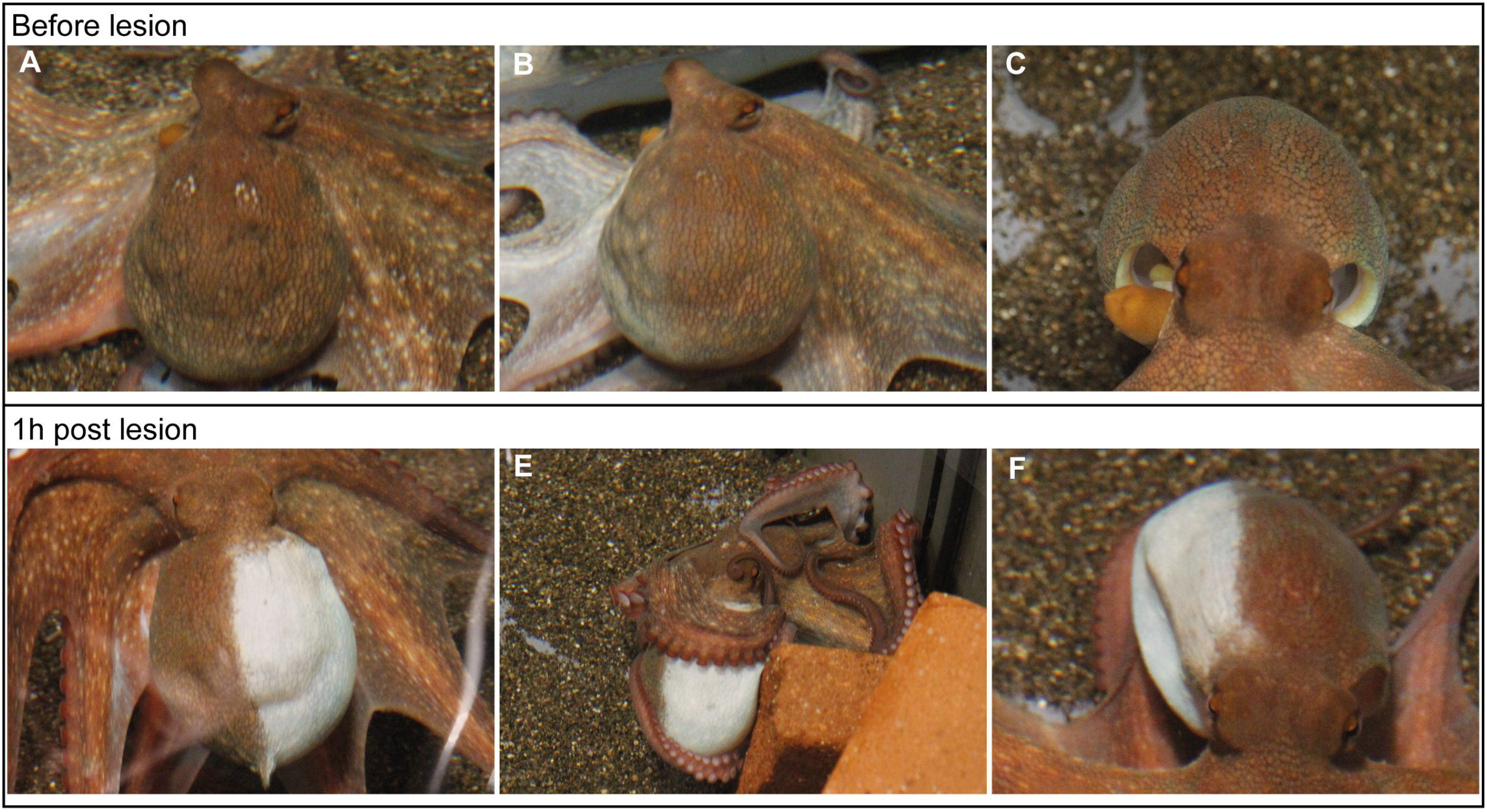
Loss of function. Uninjured octopuses are able to perform the whole range of skin patterning on both sides of the mantle bilaterally (**A**) or controlling each side unilaterally (**B**). Normal breathing is detected on both sides of the mantle through expansion and contraction of mantle openings (**C**). After surgery (one hour p.l.), complete paling of the skin (**D**) and paralysis of respiratory muscles (**F**) is detected ipsilateral to the lesion. Grooming behaviour is observed in some of the injured animals soon after lesion (**E**).

All octopuses were presented a live crab 60 min after recovery and promptly attacked it in less than 30 s performing their normal predatory behaviour; in most occasions a Full Attack response was observed (see descriptions included in: Borrelli et al., 2006; Maldonado, 1963a; Maldonado, 1963b; Packard, 1963). In order to evaluate any effect of the lesion, we considered octopuses’ predatory performances (Median Latency to attack before (bef) and after (aft) lesion, injured: LA_befInjured_ = 6.0 s, LA_aftInjured_ = 5.9 s, p = 0.966; control before and after anaesthesia: LA_befControl_ = 6.2 s, LA_aftControl_ = 6.9 s, p = 0.993; after Wilcoxon signed-rank, N = 34) and found no significant differences. We also compared predatory performances before and after anaesthesia or surgery in the two groups (LA_befInjured_ vs LA_befControl_: Z = −0.343, p = 0.731; LA_afterInjured_ vs LA_afterControl_: Z = −0.527, p = 0.598; both after Mann-Whitney U test, N_1_, N_2_ = 34), confirming the two groups behaved similarly, independently from the treatment.

Furthermore, we considered type of attack (i.e. SIDE of the approach: left or right) to assess whether octopus’ behaviour was affected by treatment. Animals approached the prey consistently as they were used to do before lesion (SIDE L vs R, injured: p = 0.109; control: p = 0.549; both after McNemar’s test, N = 34). Similarly, we tested SIDE between groups and found no significant difference (SIDE_befInjured_ vs SIDE_befControl_: χ^2^ = 0.23, p = 0.627; SIDE_afterInjured_ and SIDE_afterControl_ χ^2^ = 1.47, p = 0.224; chi-squared test after Yates correction, N = 68).

### Loss of function and recovery

Before lesion animals were fully able to exhibit their specific body pattern depending on the need, situation or stimulus they had to face in the tank and, in particular, to control bilaterally and unilaterally the chromatophores on each side of the mantle (Fig. 1 A, B; see also Supplementary Material, **Fig. S2** A, B). However, and immediately after recovery from surgery, half of the mantle (right side of the animal, i.e. the lesioned side) appeared white due to absence of any chromatic and textural pattern (Fig. 1 D). The contralateral side of the mantle retained the ability to perform the full range of body patterns, as in a normal behaving animal (Fig. 1 D).

We also observed paralysis of the mantle muscles on the lesioned side (Fig. 1 C, F).

One to two days post-surgery brown spots appeared on a pale background in the denervated area (see Supplementary Material, **Fig. S2** C) (see also Imperadore et al., 2017). At about the end of the first or at the second week post-surgery, injured octopuses exhibited some chromatic patterns on the lesioned side (see Supplementary Material, **Fig. S2** D) matching the one performed on the contralateral un-injured side (at least in tone and grooves; **Fig. S2** E). However, this capability was retained only when animals were at rest. Once moving around the tank and/or performing an attack/approach during predatory response, immediate paling of the lesioned side was observed (see Supplementary Material, **Fig. S2** F). We did not notice further improvement after two weeks post-lesion. Full reappearance of the normal colour pattern of the skin occurred (injured *O. vulgaris* both at rest and during attack) after 45 days and in some cases four months after injury (see Supplementary Material, **Fig. S2** G, H; see also bottom right picture in Fig. 2 A). We observed recovery of the ability of controlling skin texture (linked to contraction and relaxation of papillae) from about one month after lesion (data not shown).

**Figure 2.**
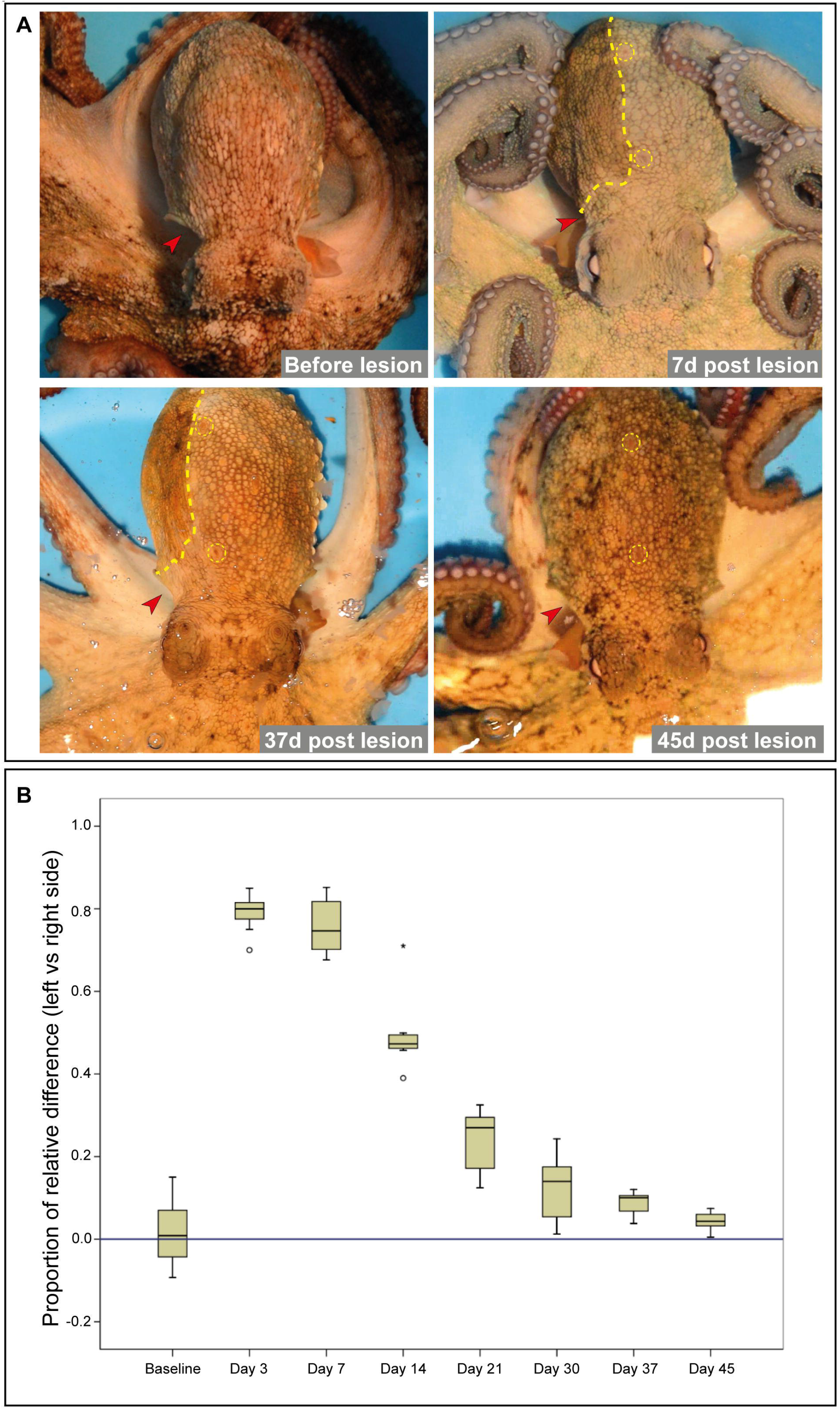
Recovery of the breathing function. (**A**) Pallial nerve transection determines interruption of the neural circuitry connecting the centralized brain to mantle muscles and chromatophores. The normal appearance of animals (as shown in upper left image) is disrupted. Seven days p.l. the mantle muscles on the lesioned side (marked by red arrowheads in all the images of the panel) are visibly paralyzed. The skin pattern on the same side of the mantle (highlighted by a yellow dotted line) is homogeneously coloured but greatly differs from the contralateral (uninjured side) and from the mantle of the animal before surgery. Restoration of the breathing function appears evident at 30-37 days p.l. and can be easily recognized by expansion and contraction of the mantle opening (red arrowhead, lower left image). No significant difference in muscles contraction was observed at 45 days p.l. but an improved control on skin pattern is detected (lower right image). Main papillae (highlighted by yellow dotted circles) are here used as skin landmarks. (**B**) Box plot showing the proportion of the relative differences of maximum extension of left (control) vs right (lesioned) side mantle openings of *O. vulgaris* before (baseline) and after lesion (N=14). Relative difference equal to zero (see reference line in the graph) is considered the normal condition. The plots show loss and recovery of breathing function with time (from day three to 45 p.l.).

Pallial nerve transection also induced impairment of mantle contractions required by normal breathing (first week p.l., Fig. 2 A, upper right), both when compared to contralateral mantle side or to the same side before lesion (Fig. 2 A, upper left). We did not notice significant progresses during the second week p.l., but improvement was observed between the third and fourth week p.l.. Overall, we observed significant departure from the baseline condition immediately after injury (Fig. 2 B) and a recovery to the normal condition starting from the fifth week after lesion. Efficient breathing on the lesioned side appeared restored only after about one month (between 37 and 45 days after surgery, respectively lower left and right in Fig. 2 A).

We observed a trend in variations in the efficiency of breathing events after lesion (contractions/expansion of the mantle openings; χ2 = 45.02, N = 7, p < 0.001, after Friedman ANOVA; Fig. 2 B). The recovery continued to progress with time, with the difference between day 30 and day 37 p.l. not resulting significant (Day 30 vs. Day 37: Z = −0.68, N = 7, p = 0.499; Monte Carlo p < 0.575, 95% CIMC = 0.565-0.585, after Wilcoxon signed-ranks test; Fig. 2 B).

### Regeneration of the pallial nerve

Backfilled contralateral control nerves did not show any differences in structure compared to naïve control nerves (see Supplementary Material, **Fig. S3** and **Video** S1). Axons were traced from the nerve to the ganglion (see Supplementary Material, **Fig. S3** Ab, Ac); inside the ganglion, fibres grow around motoneurons and form the neuropil (see Supplementary Material, **Fig. S3** Ac, left). Some fibres were seen to exit through the stellar nerves to innervate chromatophores in the skin (see Supplementary Material, **Fig. S3** Ac). We also observed fibres originating from the centripetal cells of the stellate ganglion (see Supplementary Material, **Fig. S3** Ac, **Video S1**).

Lesion of the pallial nerve determined the formation of two stumps, a central one still connected to the brain (which was used for the tracing experiments), and a peripheral one, connected to the stellate ganglion in the periphery. Within the site of lesion and a few days post-surgery, a bridge of connective tissue was formed (see Supplementary Material, **Fig. S4** Ab, Ab’). Five days p.l. fibres were not seen to cross the lesion site, but remained confined to the backfilled stump. When transected nerves were traced after 30-45 days after surgery, axons were seen to cross the gap between the lesion and the ganglion (Fig. 3 A, B; see also Supplementary Material, **Video S2**). Even though axons appeared to regenerate in several directions, the majority of the tracked fibres found the right path allowing them to enter the ganglion (Fig. 3 A, B), where they formed a network around motoneurons (Fig. 3 A, B, Bb’“). Furthermore, some of these axons were able to reach the stellate ganglion and also to leave it through the stellar nerves to reach their final targets (i.e. chromatophores). At this time point (30-45 days p.l.) no positive cells (i.e. centripetal cells) were ever observed inside the stellate ganglion as instead observed in naïve and contralateral control ganglia (see Supplementary Material, **Fig. S3** Ac, **Video S1**).

**Figure 3.**
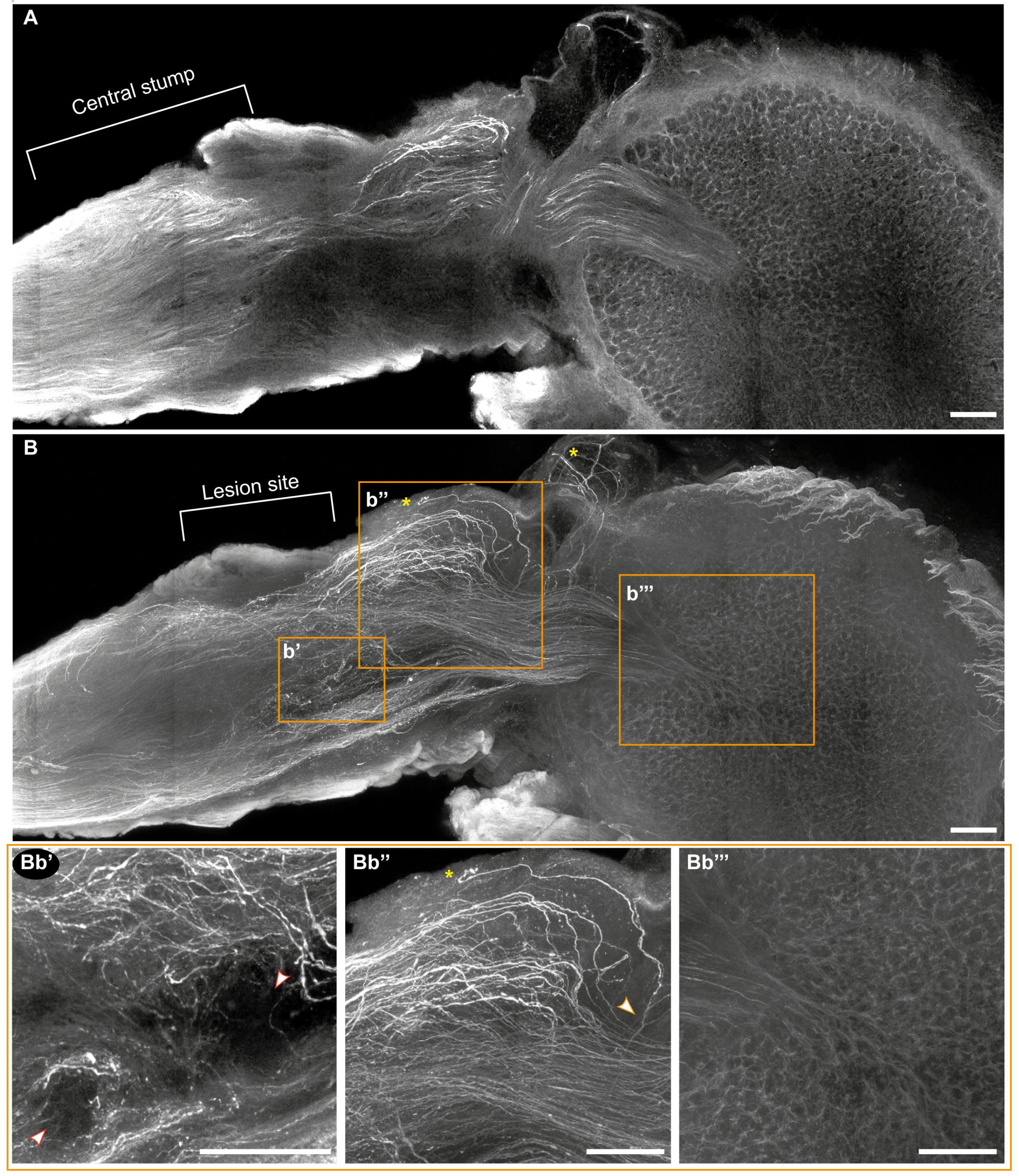
Backfilling of the nerve 45 days post lesion. After 45 days from the lesion, regenerating fibres bridge the gap between the two stumps, enter the stellate ganglion and in it they form a network among motoneurons (**A, B**). In (**A**) it is possible to observe the central stump retaining a spike-like structure with fibres showing a homogeneous calibre and a straight direction. (**B**) Proceeding deeper in the nerve in the area of lesion, regenerating fibres appear to grow in several directions even though majority of them are well-organized and directed toward the stellate ganglion. Regenerating axons show varicosities and variable calibre (**Bb’, Bb“**). In the original lesion site it is also possible to observe fibres turning around non-fluorescent obstacles (red encircled arrowheads in **Bb’**). Some fibres are also seen to deviate and grow into the musculature or backwards (yellow asterisks in **B, Bb“**). Axon branching (orange encircled arrowhead in **Bb“**) and tip swelling (yellow asterisk in **Bb“**) were also highlighted. Scale bar: 250 μm.

Regenerating fibres at the lesion site were partly well organized in bundles, with each bundle able to direct toward the ganglion. Noteworthy, some of these axons were found close to the stellate ganglion, but did not make it through and were even seen to grow backward and into the musculature (Fig. 3 B, Bb“).

The regenerated part of the growing axons did not appear to have a homogeneous calibre as in the case of intact fibres, but they showed multiple varicosities and some swelling. Branching and axon tip swelling was also observed for some of the axons that were not able to enter the ganglion (Fig. 3 Bb“).

At the level of the injury, regenerating fibres appeared to grow around non-fluorescent obstacles, which apparently diverted axons from a straight route (Fig. 3 Bb’, see also Supplementary Material, **Video S2**) possibly being the scar or debris leftovers.

Backfilling of pallial nerves at complete functional recovery showed a chaotic arrangement of fibres at the lesioned site, even though majority of axons appeared to regenerate in the correct direction, i.e. toward the stellate ganglion (Fig. 4 Ab, Nb). We detected the presence of fibre thickenings (Fig. 4 Ab, Ab’) never observed at other time points or in the control nerves; varicosities and branching were still observable (Fig. 4 Ab’). Fibres were able to reach the ipsilateral stellate ganglion, form a net around motoneurons (Fig. 4 Ac) and exit through the stellar nerves. Centripetal cells appeared traced back into the ganglion (Fig. 4 Ac).

**Figure 4.**
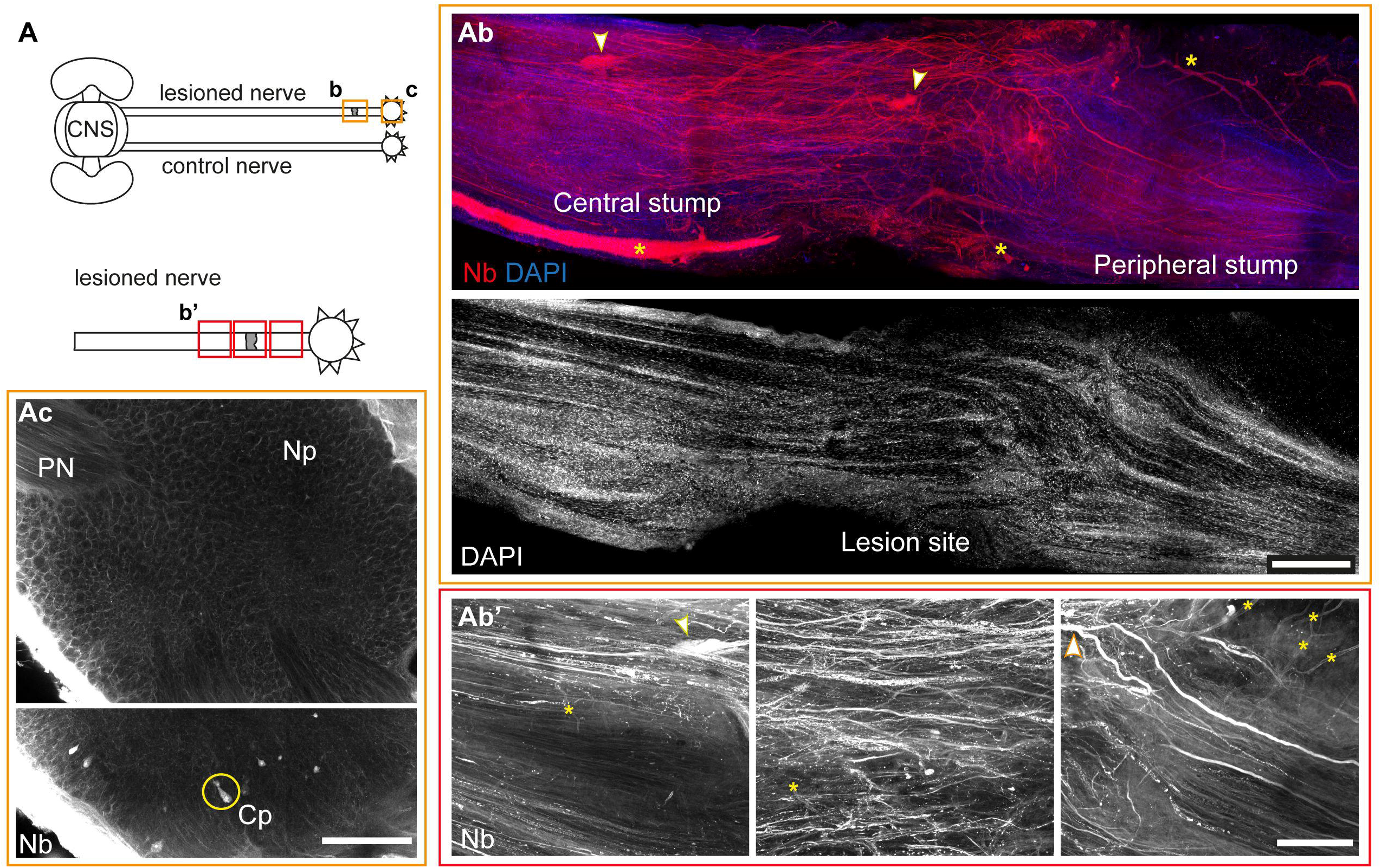
Backfilling of the nerve five months post lesion. (**A**) Schematic drawing of the connections between the central brain (CNS) and the periphery through the pallial nerves (PN). The orange rectangles highlight the areas depicted in (**Ab** and **Ac**), corresponding to the lesioned pallial nerve (**Ab**) and the ipsilateral stellate ganglion (**Ac**). The red rectangles in the schematic drawing in (bottom drawing in **A**), representing an enlargement of the lesioned pallial nerve and ipsilateral stellate ganglion, highlight the areas depicted in (**Ab’**). (**Ab**) shows an injured pallial nerve five months after surgery. Axons, traced through neurobiotin (Nb in red), were found to cross the site of lesion, connecting the two stumps. While far from injury fibres retain the classic and well-ordered appearance, in the lesion site they result more chaotic, growing in several directions. Fibre thickenings of unknown function are visible in the nerve (highlighted by yellow encircled arrowheads). Neurobiotin also highlighted an increased number of vessels and small capillaries in the lesioned nerve (yellow asterisks). DAPI counterstain (blue in the merged image) allows identification of the original site of transection, which appears as a tissue interruption. In (Ab’) enlargements of three different areas of (**Ab**) are presented, highlighting the presence of axon varicosities, fibre branching (orange encircled arrowheads) and thickenings (yellow encircled arrowheads). An increased number of vessels and capillaries is visible in these areas (yellow asterisks). (**Ac**) Fibres from the regenerating nerve reach the ipsilateral stellate ganglion, form a net around motoneurons, and exit through the stellar nerves (SN). Centripetal cells are traced back in the ganglion (yellow circle). Scale bars: (**Ab, Ac**) 500 μm; (**Ab’**) 150 μm. Abbreviations: CNS, central nervous system; Cp, centripetal cells; Nb, Neurobiotin; PN, pallial nerve; Np, neuropil.

Tracing experiments also highlighted an increased number of vessels and small capillaries in lesioned nerves (Fig. 4 Ab, Ab’) again not visible in control nerves, suggesting the occurrence of angiogenesis during the regenerative events.

The site of lesion resulted recognizable as a tissue interruption (Fig. 4 Ab, DAPI) being more evident on the dorsal side while getting less evident in the middle and ventral sides.

### Functionality of regenerated axons in the pallial nerve

Regenerated fibres in the pallial nerve express axonal properties, as tested through two bipolar electrodes placed before the lesion site (central stump) and close to the stellate ganglion (peripheral stump) on the surgically isolated pallial nerve of regenerated animals five months p.l. (Fig. 5 A). Electrical stimulation via the stimulating electrode close in the central stump was found to systematically induce activity in axons crossing the lesion site in the recording electrodes close to the stellate ganglion in all experiments (Fig. 5 B). With increasing stimulation amplitude, the extracellular recorded compound action potential increased in two out of three preparations. Such increase in compound action potential amplitude reveals that axons of different diameters are recruited upon increasing stimulation strength (Fig. 5 C). Interestingly, the activation of axons upon electrical stimulation in the contralateral control pallial nerve (Fig. 5 A, C), showed a similar systematic correlation between stimulation strength and amplitude of the compound action potential.

**Figure 5.**
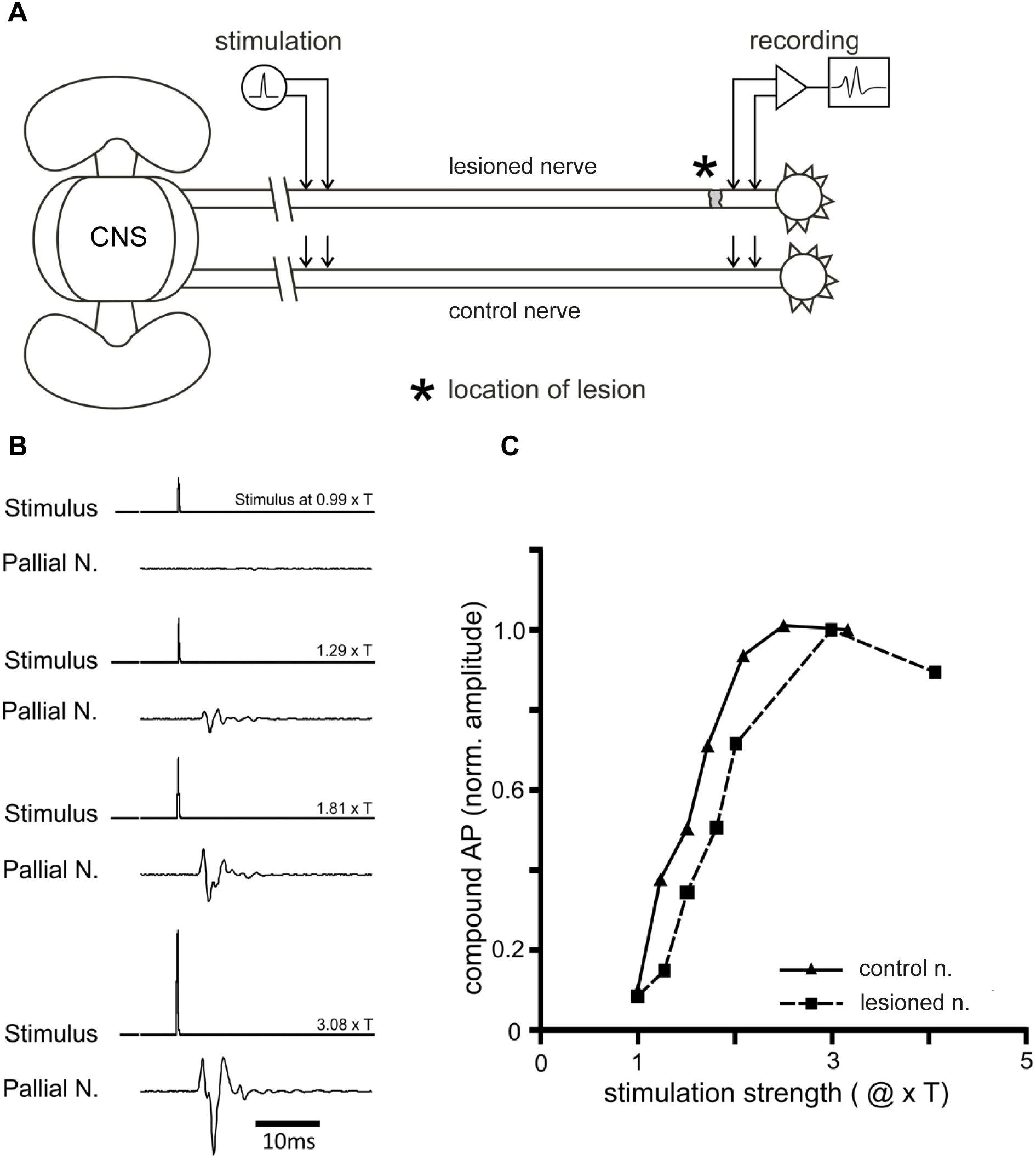
Effect of the electrophysiological stimulation of axons in the pallial nerve. (**A**) Schematic depiction of the stimulation setup. Bipolar extracellular electrodes were placed on two locations of the surgically isolated pallial nerve, which connects the brain and the stellate ganglion both for the regenerated pallial nerve (top) and the control condition (bottom). (**B**) Effect of electrical stimulation on the regenerated pallial nerve at four stimulation intensities in one preparation: 0.99xT (below threshold) and 1.29, 1.81, 3.08 (above threshold). Please note that the compound action potential increases with increasing stimulation amplitude (see text). (**C**) Dependence of the normalized amplitude of the compound action potential induced by electrical stimulation in one regenerated nerve as compared to a control nerve. Please note that the values plotted are means (n=5 stimulations each). Depiction of standard deviation is not visible as the amplitude values of the compound action potentials upon electrical stimulation merely differed between trials for a given stimulation strength.

## Discussion

Lesion of one of the two pallial nerves in *O. vulgaris* leads to the interruption of the neural circuits between the central brain and the stellate ganglion in the mantle (i.e. periphery). As each nerve and related ganglion control only one side of the mantle (see Supplementary Material, **Fig. S1**), loss of function can be immediately recognized on the ipsilateral side of the injury (see Fig. 1): this corresponds to skin paling (as originally observed by Fredericq, 1878) due to lack of control of chromatophores in the skin, and to the impairment of muscle function for the control of mantle breathing movements and papillae. The contralateral side preserves intact functionality (Imperadore et al., 2017). These dramatic changes appear reversible, due to the ability of the pallial nerve in functionally regenerating (Sanders and Young, 1974; Sereni and Young, 1932).

We evaluated timing and pathways of neural fibre regeneration and functional recovery for this nerve after complete transection.

All octopuses recovered the ability in controlling skin papillae at around one month following surgery. First signs of mantle muscle contraction recovery, described as muscular tone improvement in Imperadore et al. (2017), were observed already at 14 days p.l.. Breathing recovery was resumed between 30-37 days post-surgery for all the individuals.

These findings are also consistent with the morphological data obtained from the imaging of the backfilled pallial nerves. Axons from the central stump, indeed, required a comparable amount of time to reach motoneurons in the stellate ganglion (controlling breathing muscles; Monsell, 1977; Young, 1972), forming a net around them and realistically building synapsis.

The time required by fibres to reach the chromatophores in the skin was consistently longer, as fibres had to reach the ganglion and exit from through the stellar nerves thus to travel for several centimetres to innervate the millions of chromatophores spread in the skin of the mantle (estimated around 200 per mm2 in O. vulgaris: Messenger, 2001). All octopuses kept alive for five months after lesion were able to fully recover skin patterning. However, for one of them 45 days were sufficient, while two required around two months and the last two required four months for complete recovery. In all these cases, anaesthetized animals (for human killing) did not show wandering clouds on the skin of the lesioned side as observed at previous time points. The skin appeared as pale and unreactive to pinch as the contralateral side of the mantle and comparable to what observed in uninjured animals under anaesthesia.

If we consider regeneration and re-innervation in terms of fibre speed growth, we can speculate that axons from the central stump reach the stellate ganglion and completely re-establish synapsis with motoneurons at a speed of 10 μm per hour, as they had to cover a distance of ∼3.5 mm to reach the ganglion and the same length to pass through the ganglion. Our data confirm the rate of growth originally described by Sereni and Young (1932).

However, fibres directed from the nerve to the chromatophores apparently regenerate faster. Fibre speed in this case appears to be approximatively between 34 and 90 μm per hour, considering 7 mm distance to exit the ganglion and around 90 mm to reach the farther chromatophores in the mantle, as it is the case at complete recovery.

The large number of fibres found in the lesioned area together with their intricate pathways made it difficult to assess the true origin of fibres. Even though we tracked axons from the central stump, we are not able to assert with certainty that these fibres were originating from the central brain and not from the peripheral stump of the pallial nerve, which also start to regenerate a few days post injury (Imperadore et al., 2017).

Fibre varicosities, swelling, branching, obstacle deviation and thickenings were visible in the regenerating nerves; the latter were never observed in uninjured or contralateral nerves. Sereni and Young (1932), also report regenerating fibres deviating around obstacles; these may represent residue of scars or debris of the degenerating materials, both identified few days p.l. (Imperadore et al., 2017; 2018) and also described to occur in mammals during peripheral nerve regeneration (Kang and Lichtman, 2013).

Swelling, branching and fibre aberration appear to persist for months; the regenerated nerves never retrieved their original appearance, not even five months p.l., although all animals had perfectly recovered the altered functions by this time.

We cannot exclude that alternative mechanisms might also be involved and aid to chromatophore re-innervation, such as intervention through collateral nerve sprouting of the stellar nerves from the opposite stellate ganglion (as suggested by Packard, 1995). This is further supported by the fact that the mantle midline and the areas close to it return functional much earlier than other areas (see Supplementary Material, **Fig. S5**). Further investigation is required to support this hypothesis.

Our electrophysiological experiments on samples taken five months p.l. confirmed that lesioned nerves are able to re-establish functional connection between the brain and the periphery. The regenerated fibres express axonal properties, including voltage-dependent ion channels, generate and propagate action potentials. Electrical stimulation via the stimulating electrode close to the central stump was found to systematically induce activity in axons crossing the lesion site in the recording electrodes close to the stellate ganglion. Increasing stimulation amplitude resulted in an increase in the extracellular recorded compound action potential. Such increase in compound action potential amplitude reveals that axons of different diameters are recruited upon increasing stimulation strength (Büschges et al., 1992; Hooper and Schmidt, 2017).

Previous studies on regeneration of the pallial nerve in the octopus (Sanders and Young, 1974; Sereni and Young, 1932) considered 60 to 150 days as required for complete recovery of function of body patterning after injury (in the original study either through crush or cut). According to the Authors and in some cases this never occurred and the animal continued to show lack of full recovery (Sanders and Young, 1974; Sereni and Young, 1932). Crushing of the nerve required nine to ten weeks for the complete recovery of body patterns. However, no octopuses showed signs of colour pattern recovery until 50 days p.l., neither in summer nor in autumn when experiments were carried out (Sanders and Young, 1974; Sereni and Young, 1932). In the original study, six out of 10 animals recovered complete colour pattern (between 60 and 69 days), while only for two out of 10 octopuses it was possible to observe functional recovery of textural components (i.e. papillae; between 30 and 50 days). On the other hand, only four animals (out of 10) undergoing complete transection of the pallial nerve recovered colour patterns (signs of recovery already at 30 days). In this case the complete regain of function required 109 days. In addition, seven octopuses (out of 10) recovered the ability in raising papillae (Sanders and Young, 1974; Sereni and Young, 1932).

Strangely enough, when regenerated nerves were inspected, the authors could not determine any clear correlation between histological appearance and functional recovery, also stating the inability in following fibres pathways and orientations (Sanders and Young, 1974; Sereni and Young, 1932).

In summary, our results show that during regeneration neural connectivity between the brain and the stellate ganglion can be re-established by the formation of axons crossing the lesion site.

In our view the octopus pallial nerve provides an exceptional example of regeneration in cephalopods, because: ***i.*** regeneration is always relatively quick and efficient; ***ii.*** it is immediately possible to evaluate loss and regain of function by evaluating respiration (i.e. breathing movements) and skin patterning on the mantle of the animal; ***iii.*** each animal has a pair of nerves, one on each side of the mantle, allowing the same animal to serve as experimental and control; ***iv.*** the same nerve can be lesioned several times providing almost complete regeneration at each instance (Packard, 1995); ***v.*** the majority of the cell soma of axons in the pallial nerve resides inside the brain, but others in the ganglia at periphery. Both can regenerate, allowing a comparison of regeneration ability of the two ‘systems’, one being spatially more centrally the other more peripherally located; ***vi.*** *Octopus* represents a “simpler” animal compared to vertebrates. However, regeneration/degeneration phenomena following lesion are found similar to those happening in higher vertebrates (Wallerian degeneration).

## Acknowledgements

Authors are thankful for the assistance and guidance given by Dr Astrid Schauss and Dr Christian Jüngst (CECAD Imaging facility, CECAD Research Center, Cologne, Germany) for accessing the imaging facility and for the use of Leica SP8 multiphoton microscope. Authors are also grateful to Leica Microsystems (P. Romano, K. Orellana) for assistance during various phases of this work. This work benefited of the networking initiative of the COST Action FA1301 – Cephs*In*Action.

## Competing interests

No competing interests declared

## Authors’ contributions

PI carried out experiments, analyzed the data, processed images and drafted the manuscript; MD and AB performed electrophysiology experiments; AB contributed to the experimental design and manuscript editing; DP and AO assisted in the image acquisition and processing from thick samples; GF supervised the work, designed the experiments and revised the final manuscript. All authors contributed to the final writing of the manuscript and approved the final article.

## Funding

This work was supported by the Stazione Zoologica Anton Dohrn and RITMARE (Flagship Project - MIUR & SZN) to GF, and by the Association for Cephalopod Research - CephRes to PI. This work also benefited of the infrastructure part of the project “Potenziamento di una piattaforma integrata per lo studio di malattie umane di grande impatto attraverso l’uso del system phenotyping di modelli animali: Mouse e Zebrafish clinic (MouZeCLINIC)” to the SZN (PONREC PONa3_00239).

